# Prevalence and genotypes of *Chlamydia psittaci* in pet birds of Hong Kong

**DOI:** 10.1101/2024.02.19.581005

**Authors:** Jackie Cheuk Kei Ko, Yannes Wai Yan Choi, Emily Shui Kei Poon, Nicole Wyre, Jennifer Le Lin Go, Leo Lit Man Poon, Simon Yung Wa Sin

## Abstract

Psittacosis, or parrot fever, is a zoonotic disease caused by *Chlamydia* species associated with birds. One of the causative agents of the disease is *Chlamydia psittaci*, which is often carried by psittacine and can be highly pathogenic and virulent to humans. In Hong Kong, a city with high population density, psittacosis is a notifiable disease with over 60% of cases in the last decade resulting in hospitalization. However, the source of transmission of *C. psittaci* and its prevalence in pet birds in Hong Kong are currently unknown. To evaluate the risks of psittacosis transmission through pet birds, we tested the presence of *C. psittaci* and determined its genotypes in samples obtained from 516 captive birds from households, pet shops, and a veterinary hospital in Hong Kong. Results revealed that five samples (0.97%), collected from budgerigars and cockatiels, were *C. psittaci*-positive, while 80% of them were obtained from pet shops. The identified strains belonged to genotype A and were closely related to the strain SP15 that was previously discovered in mainland China. Our study highlights the need for genotyping *Chlamydia* species in psittacosis patients to understand their source(s) of origin.

## Introduction

Psittacosis, also known as parrot fever, is a zoonotic disease caused by avian-associated *Chlamydia* species, with *Chlamydia psittaci* being the major and most studied causative agent (Balsamo et al. 2017). In humans, *C. psittaci* infection can be fulminant or subclinical (Zhang et al. 2022), leading to influenza-like illnesses, pneumonia, and even death, which is especially common in the elderly (Billington 2005; Tolba et al. 2019). Although antibiotic treatment for psittacosis is available, the disease remains an important health concern, especially for immunosuppressed individuals (Billington 2005).

The host range of *C. psittaci* is broad. Over 465 bird species in 30 orders have been found vulnerable to *C. psittaci* infection (Harkinezhad et al. 2009). While birds are the primary carriers, *C. psittaci* has been found in various animals, including mammals, e.g., dogs, cats, and horses, as well as reptiles, e.g., crocodiles, lizards, and tortoises (Hogerwerf et al. 2020; Inchuai et al. 2020). To date, 16 genotypes of *C. psittaci* have been discovered based on sequences of the outer membrane protein A (*OmpA*) gene (Radomski et al. 2016), and each genotype has differential host preferences (Beeckman et al. 2009). Genotype A is associated with psittacine birds and has the highest virulence and pathogenicity in humans (Harkinezhad et al. 2009; Heddema et al. 2015). Other genotypes have been isolated from waterfowls, chickens, turkeys, pigeons, passerines, and other avian or mammalian species (Liu et al. 2019; Vanrompay 2019). According to Sukon et al., the global prevalence of *C. psittaci* is highest in Galliformes and Columbiformes (Sukon et al. 2021), but human infections from these genotypes are rare due to their low virulence in humans (Sukon et al. 2021). Nonetheless, all *C. psittaci* genotypes can cause diseases and are transmissible to humans (Beeckman et al. 2009).

Surveys worldwide have monitored *C. psittaci* prevalence and genotypes in captive, domestic, and wild birds since the 1980s (Sukon et al. 2021). Captive psittacine birds, particularly pet parrots, raise concerns, as they were reported to cause the most psittacosis outbreaks (Hogerwerf et al. 2020; Kozuki et al. 2020). However, data on the prevalence and genotypes of *C. psittaci* in captive psittacines in Asia are limited. In the last decade, detection, and characterization of *C. psittaci* in pet parrots have only been conducted in a few Asian countries or regions, with prevalences ranging from 3.1% to 20.65% (Sukon et al. 2021). Given the danger posed by psittacine-carried strains and the ongoing public health concern in many parts of Asia (Yin et al. 2015; Kozuki et al. 2020; Abd El-Ghany 2020; Shi et al. 2021; Lee et al. 2023), understanding the epidemiology of *C. psittaci* in pet parrots is crucial.

Noteworthily, psittacosis has been a persistent public health concern in Hong Kong, with at least ten cases reported annually in the past decade. It was designated as a notifiable infectious disease under the Prevention and Control of Disease Ordinance in 2018 (CHP 2020). Among recorded cases in the last decade, a quarter reported contact with pet birds, primarily parrots, or their droppings. Over 60% of patients were hospitalized, with two deaths occurring due to respiratory failure or pneumonia (CHP 2020). Despite the recurring cases and the high-density pet bird population in Hong Kong (ADM capital foundation 2022), there is a lack of information regarding the source of transmission, prevalence, and genotypes of *C. psittaci* in birds and humans. Assessing infection rates, distribution, and genotypes of *C. psittaci* in pet birds, especially parrots, and their owners is essential for understanding the epidemiology of the disease in the community.

The purpose of the study reported here is to determine the prevalence and genotypes of *C. psittaci* in captive psittacine and passerine birds from pet shops, households, and a veterinary clinic in Hong Kong. A specific nested PCR assay targeting the *OmpA* gene was developed to detect *C. psittaci* DNA, and genotypes were identified through *OmpA* gene sequencing. These findings will be valuable for preventing and managing psittacosis transmission in Hong Kong and expanding our knowledge about *C. psittaci* carried by pet birds in Asia.

## Materials and Methods

### Sample Collection

Between November 2019 and January 2022, a total of 516 fecal samples were collected (Supp 1). These samples were obtained from the cages of captive birds from 218 households (N=346), 4 pet shops (N=54), and a veterinary hospital (N=116). The samples encompassed a wide range of popular pet bird species, including 43 psittacine species and 7 passerine species. Most samples were collected from individual birds, except for 17 samples from pet shops, which were collected from 4 cages housing multiple budgerigars or cockatiels. During the sampling process at the veterinary hospital, 8 of the birds were receiving antibiotics at the time. Among the antibiotics used were doxycycline, which had been added to the drinking water of a common hill myna (*Gracula reliqiosa*) a week prior to sampling, and enroflaxacin, which had been administered to treat a grey parrot (*Psittacus erithacus*), a peach-faced lovebird (*Agapornis roseicollis*), and a Pacific parrotlet (*Forpus coelestis*) (Lindenstruth et al. 1993; Butaye et al. 1997; Failing et al. 2006). Other antibiotics used were generally not effective against *C. psittaci*, including chloramphenicol, amoxicillin-clavulanic acid, and metronidazole. All samples were stored in 100% ethanol at -20°C shortly after collection to preserve DNA in the samples. Whenever possible, information on the age, sex, symptoms, and medical history of each sampled bird, was obtained.

### DNA Extraction

For DNA extraction from bird fecal samples, the E.Z.N.A. Stool DNA Kit (Omega Bio-tek, Norcross, USA) was used. Approximately 200 mg of each sample was used for extraction following the manufacturer’s protocol. Sample homogenization was achieved using a TissueLyser II (Qiagen, Hilden, Germany) with 5mm stainless steel beads (Qiagen). The eluted DNA was stored in 30-50μL of elution buffer at -20°C.

### Nested PCR Assays for Detecting C. psittaci

Nested PCR assays were developed to detect *C. psittaci* DNA in bird fecal samples. Using the *OmpA* sequences of *C. psittaci*, *Chlamydia trachomatis*, and *Chlamydia pneumonia* retrieved from GenBank, two pairs of primers were designed to amplify two different regions of the *OmpA* gene of *C. psittaci*, respectively (Supp 2). Primer pairs O1 and N1 were used to amplify the variable domain (VD) I, while pairs O2 and N2 were used to amplify VD III-IV.

In each reaction for the first PCR, 5μL of extracted DNA was used as template in a total volume of 25μL, with primer concentration of 0.6 μM (IDT, Coralville, USA). On the other hand, reaction mixture for the nested PCR consisted of 1μL of amplified first PCR product as template in a total volume of 25μL, with primer concentration of 0.6 μM (IDT). Touchdown conditions were used for the first PCR, while conventional PCR condition was used for nested reactions. The temperature condition for the first PCR was 95°C for 2 min, followed by 40 cycles of 95°C for 30 sec, 63.5-59.5°C (63.5-60.5°C for the first 4 cycles and 59.5°C for the remaining 36 cycles) for 30 sec, 72°C for 45 sec, and 72°C for 5 min. For the nested PCR, the condition was 95°C for 2 min, followed by 40 cycles of 95°C for 30 sec, 59.5°C for 30 sec, 72°C for 45 sec, and 72°C for 5 min.

The study employed Amplirun *C. psittaci* DNA Control (Vircell Microbiologists, Granada, Spain) as a ten-fold diluted positive control and UltraPure water (Invitrogen) as a negative control in the initial PCR reactions. To avoid potential cross-contamination between samples, 28 samples were tested per batch, along with one positive and one negative control. In total, 30 rounds of detection PCR were conducted to analyze all samples. The expected results were obtained in all positive and negative control reactions, which were confirmed by DNA gel analysis. All PCR products with expected size were sequenced by BGI (BGI Genomics, Hong Kong), and the sequences were analyzed using Geneious Prime 8.1.9. The identities of sequences were verified by performing searches in the basic local alignment search tool (BLAST) against GenBank (NCBI) database.

### *OmpA* Gene Amplification for Genotype Identification

*OmpA* gene amplification was performed to identify genotypes of *C. psittaci* in positive samples. Initially, the protocol described by Madani et al. was attempted but yielded suboptimal amplification (Madani et al. 2013). Therefore, three pairs of primers were designed based on *C. psittaci OmpA* sequences from GenBank, to amplify three overlapping regions within the gene, to obtain the full *OmpA* sequences. (Supp 3). Primer pairs C2 and C3 were used for VD I-II and VD III-IV amplification, respectively, while primer pair C3 was used to amplify VD I-IV. The reaction mixtures consisted of 5μL of extract-ed DNA as a template in a total volume of 40μL, with primer concentration of 0.4μM (IDT). Touchdown conditions were used for the reaction. For reaction C1, the temperature condition was 95°C for 2 min, followed by 40 cycles of 95°C for 30 sec, 62.5-60.5°C (62.5-61.5°C for the first 2 cycles, and 60.5°C for the remaining 38 cycles) for 30 sec, 72°C for 45 sec, and 72°C for 5 min. For reaction C2, temperature condition was 95°C for 2 min, followed by 40 cycles of 95°C for 30 sec, 59-57°C (59-58°C for the first 2 cycles, and 57°C for the remaining 38 cycles) for 30 sec, 72°C for 45 sec, and 72°C for 5 min. For re-action C3, temperature condition was 95°C for 2 min, followed by 40 cycles of 95°C for 30 sec, 60.5-52.5°C (60.5-53.5°C for the first 8 cycles, and 52.5°C for the remaining 32 cycles) for 30 sec, 72°C for 45 sec, and 72°C for 5 min. Positive and negative controls were included using diluted *C. psittaci* DNA control and UltraPure DNase/RNase-free distilled water (Invitrogen), respectively. PCR products were sequenced and analyzed using Geneious Prime 8.1.9, with sequence identities verified through BLAST searches against GenBank (NCBI) database. Sequences ranging from 729 to 1128bp were obtained from positive samples (GenBank accession number: OP594252-OP594256).

### Statistical analyses

Confidence intervals of *C. psittaci* prevalences in each sampled species was calculated using the Reiczigel method (Lang et al. 2014). Fischer’s’ exact test was subsequently employed to test for significant differences between prevalences between sources and host species.

### Genotype Determination and Phylogenetic Relationship Construction

To determine genotypes and establish phylogenetic relationships, the *OmpA* sequences from the samples and selected reference strains were aligned using MAFFT. The reference strains (Supp 4) included those identified by Sachse et al. (Sachse et al. 2014), as well as the strain SP15 (GenBank sequence ID: EU856035) identified in Yunnan and other parts of mainland China (Zhang et al. 2015). The genotypes and strains were determined based on nucleotide similarity and the construction of a phylogenetic tree. Nucleotide p-distances were calculated using Geneious. Model selection and phylogenetic tree reconstruction were conducted using IQ-Tree (Nguyen, 2015; Trifinopoulos et al. 2016; Kalyaanamoorthy et al. 2017; Hoang et al. 2018;). A maximum likelihood (ML) tree was constructed with 1000 bootstrap replicates. The HKY+F+G4 model (Hasegawa-Kishino-Yano model with a consideration of empirical base frequencies and application of gamma rate heterogeneity) was selected (Hasegawa et al. 1985). The phylogeny was visualized using Figtree 1.4.4, with *C. abortus OmpA* sequence (GenBank: AJ004873.1) was added as the outgroup.

## Results

### Prevalence of C. psittaci in Sampled Birds

Out of 516 bird fecal samples, five (0.97%) were detected positive for *C. psittaci* (Table 1). The positive samples were collected from two parrot species, with four from budgerigars (*Melopsittacus undulatus*; 13.8%) and one from cockatiels (*Nymphicus hollandicus*; 1.61%). Fischer’s exact test did not reveal a significant difference in *C. psittaci* prevalences between host species (*P*=0.32). One of the positive samples was collected from a singly housed budgerigar from a household, while the other three were collected from a cage containing 10 budgerigars in a pet shop. It was uncertain which individual and how many individuals were infected with *C. psittaci* in this cage, since two negative samples were collected from the same cage at the same time, suggesting that at least one bird in that cage might be uninfected. The positive sample from cockatiels was collected from a cage housing seven birds of the same species in the same pet shop, and two other samples collected from the same cage were negative for *C. psittaci*. The prevalence of *C. psittaci* was found to be 7.41% in the samples collected from pet shops. This was much higher than the prevalence in households (0.29%) and the animal clinic (0.00%). The differences in prevalences among these sources were statistically significant as tested using Fischer’s exact test (*P*<0.001).

**Table 1.**
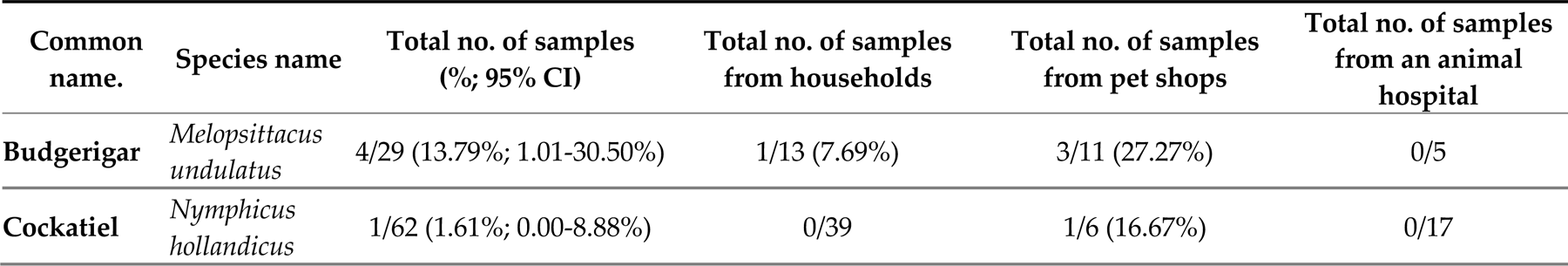
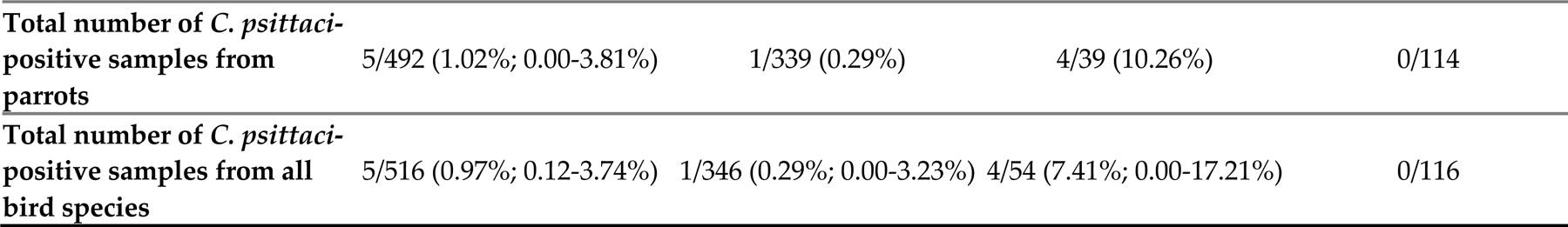
Summary of the prevalences of *C. psittaci* in bird samples from households, pet shops, and a veterinary hospital. Confidence intervals (CI) were calculated using the Reiczigel method (Lang et al. 2014).

### Genotypes of Identified C. psittaci in Bird Samples

Based on nucleotide similarity and phylogenetic analysis (Figure 1 and Supp 5), all *C. psittaci* detected in bird fecal samples were identified as genotype A. Nucleotide p-distances show that our sequences were highly similar to the strain SP15 (99.82%-100%; Supp 5), with two sequences (OP594254 & OP594256) amplified from two samples being identical to SP15.

**Fig 1.**
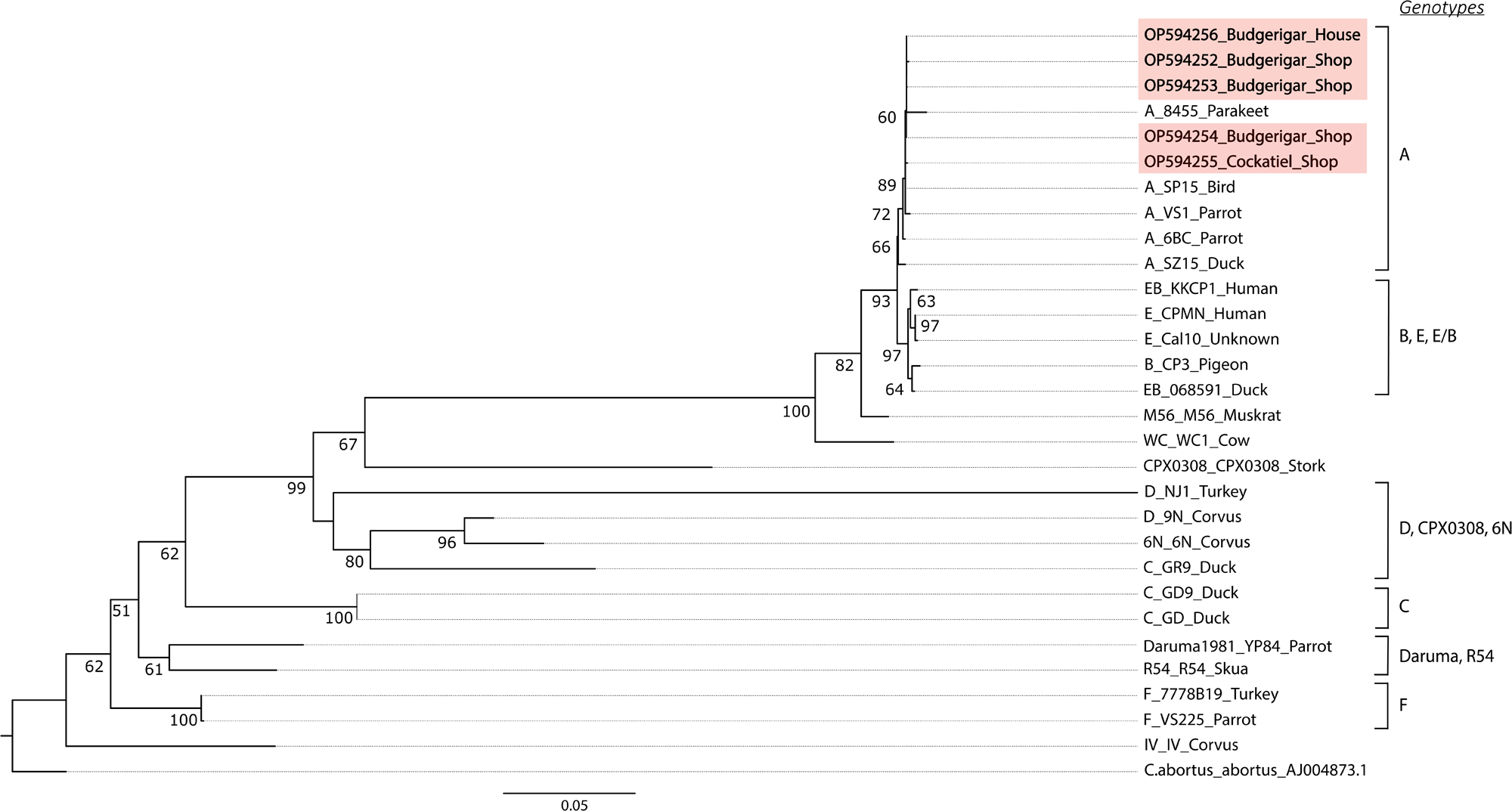
Phylogenetic tree reconstructed using 1122 bp *Chlamydia psittaci OmpA* gene sequences from positive samples in this study (shaded in light red) and reference sequences from GenBank. Bootstrap values greater than 50 are displayed at each node. *Chlamydia abortus* sequence (AJ004873.1) was included as an outgroup. Substitutions per site are indicated by the scale at the bottom.

All *OmpA* sequences amplified were highly similar to each other, with nucleotide distances ranging from 99.48% to 99.91% (Supp 6). Phylogenetic analysis reveals that the *C. psittaci* sequences in our samples formed a highly supported clade with genotype A, strains SP15, VS1, and 84-55 (Fig. 1). All genotype A sequences formed a monophyletic clade, which was sister to the clade of genotypes E and B.

## Discussion

Our study conducted the first screening of *C. psittaci* in local captive birds in Hong Kong. Despite the high-density pet bird population and active pet bird trade market (Poon 2018; Chan et al. 2021; ADM capital foundation 2022), the prevalence of *C. psittaci* (i.e., 0.97%) was lower than that of most nearby regions, including Yunnan province, Gansu province, Beijing city, and Weifang city of China. These regions reported *C. psittaci* prevalences ranging from 10.8% to 35.57% in pet birds from pet markets, zoos, or unknown sources during 2015-2016 (Cong et al. 2014; Zhang et al. 2015; Feng et al. 2016). In Taiwan, a prevalence of 3.1% was reported in breeding facilities, a bird imports corporate, a zoo, and a veterinary hospital in 2019 (Liu et al. 2019).

Among the 516 samples obtained from 50 psittacine and passerine species from households, pet shops and an animal clinic, positive samples were found in budgerigars and cockatiels, which are popular pet parrots both locally and globally. High prevalences of *C. psittaci* in these two parrot species have been reported in multiple regions or countries (Beeckman et al. 2009; Yin et al. 2015; Mina et al. 2019; Tolba et al. 2019; Lee et al. 2023), and it was suggested that *C. psittaci* is more prevalent in these two species than other parrots (Smith et al. 2005).

Notably, we observed low prevalences in several parrot species that are heavily traded, including peach-faced lovebirds, grey parrots, monk parakeets (*Myiopsitta monachus*), turquoise-fronted amazons (*Amazona aestiva*), and others. Particularly, despite sampling over 10 individuals from pet shops, we found zero prevalence in peach-faced lovebirds. This contrasts with previous studies that reported high infection rates of *C. psittaci* in parrots such as Amazons, conures, Senegal parrots, and lovebirds (Fudge 1997; De Freitas Raso et al. 2006; Feng et al. 2016; Lee et al. 2023). Similarly, we did not detect any *C. psittaci* DNA in passerine birds. Apart from occasional positive cases in finches, sparrows and mynahs, chlamydiosis has historically been a rare issue in passerine species when compared to Psittacines or Columbiformes (Andersen et al. 2000).

Among the three sample sources, the number of positive samples collected from pet shops was significantly more than the other sources (households or animal clinic), consistent with findings from previous studies that reported pet shops or breeding facilities as the primary sources of *C. psittaci* transmission (Feng et al. 2016; Tolba et al. 2019; Vanrompay 2019). Risk factors associated with pet shops, such as poor hygiene conditions, high bird density, or increased stress levels, can contribute to the proliferation and transmission of *C. psittaci* (Harkinezhad et al. 2009).

Meanwhile, based on nucleotide similarity and phylogenetic analyses results, all the identified strains belong to genotype A, which is closely associated with parrot hosts (Radomski et al. 2016). The *OmpA* sequences of these strains were highly similar to the strain SP15 found in Yunnan in 2008 (Liu et al. 2008). This strain was later detected in pet parrots from bird markets in Weifang and Beijing (Sachse et al. 2014). As they are major pet parrot production sites in China, with high *C. psittaci* prevalences, it was suggested that the SP15 strain could be disseminated from these cities to other parts of the country through parrot trade (Sachse et al. 2014). Since mainland China is a main supplier of pet parrots to Hong Kong (Poon 2018), it is possible that the detected *C. psittaci* strains originated from mainland China. Additionally, it was suggested that a majority of pet birds sourced from mainland China to Hong Kong were likely imported illegally (Cong et al. 2014), bypassing mandatory veterinary assessments and clinical observations. including those specific to Chlamydiosis (Agriculture, Fisheries and Conservation Department 2024). Such trafficked birds pose a higher risk of *Chlamydia* infection, as demonstrated in previous studies (Harkinezhad et al. 2009; Vilela et al. 2020; Ruiz-Laiton et al. 2022). Investigating the role of pet bird trafficking in the dissemination of *C. psittaci* in Hong Kong is therefore crucial.

While the presence of *C. psittaci* in pet parrots from households and pet shops indicates their potential as sources of psittacosis in Hong Kong, the exact role of pet birds in psittacosis transmission remains unclear. Three quarters of psittacosis patients in the past decade did not report any contact with birds or their secretions (CHP 2020). It is possible that certain strains can transmit the disease without relying on avian hosts, instead spreading through environmental contamination or human-to-human contact, as observed in multiple psittacosis outbreaks (Branley et al. 2016; Zhang et al., 2022). To gain a better understanding of the sources and transmission routes of *C. psittaci*, it is recommended to focus on genotyping *C. psittaci* strains in psittacosis patients at the time of diagnosis. Besides, conducting mass-scale screening of *C. psittaci* in asymptomatic humans who are in close contact with birds would be valuable. Improved knowledge of *C. psittaci* epidemiology, including their modes of transmission and host-strain associations, is essential for effective measures against psittacosis in Hong Kong and other affected regions.

In addition to *C. psittaci*, there has been a growing number of psittacosis cases worldwide caused by other *Chlamydia* species, especially *C. avium* and *C. abortus*, which are carried by pet birds (Opota et al. 2015; Liu et al. 2019). Although limited reports have described these *Chlamydia* species in parrot hosts within Asia (Chahota et al. 2006), it is crucial to conduct future research on pan-*Chlamydia* detection in pet birds of Hong Kong and other parts of Asia to under-stand their roles in causing psittacosis in regions where the disease is a notifiable problem.

Our study has a few limitations. Firstly, we only tested fecal samples, which may have resulted in a biased detection rate compared to other sampling sites. It has been suggested that fecal samples, along with cloacal samples, may have a lower positivity rate than pharyngeal swab samples (Anderson et al. 1996). Secondly, despite that the nested PCR approach had been widely used and supported by past literature (Harkinezhad et al. 2007; Madani et al. 2013; Zhang et al. 2022), a systematic comparison in sensitivity and specificity between our designed assay and other detection methods, such as qPCR and microarray-based approaches, is lacking. Our results therefore may not be directly comparable to those of other detection methods. Additionally, since this study did not include analyses of *C. psittaci* infectivity and vitality analyses, it remains unknown whether the detected *C. psittaci* DNA reflects the presence of live, transmittable bacteria. Without this information, it is difficult to evaluate the clinical and epidemiological implications of the observed *C. psittaci* DNA positivity. Further research is necessary to elucidate the transmission dynamics and clinical significance of *C. psittaci* infections in parrots in Hong Kong.

## Conclusions

The study aimed to investigate the prevalence and the genotypes of *C. psittaci* among pet birds in Hong Kong, including those from households, pet shops, and a veterinary clinic. Out of the 516 samples tested, *C. psittaci* DNA was detected in 5 cases (0.97%). All identified *C. psittaci OmpA* sequences belong to genotype A and closely resemble the SP15 strain previously found in Yunnan, Weifang, and Beijing. Notably, a significant number of positive samples were obtained from parrots in pet shops, suggesting the potential for widespread transmission of *C. psittaci* in pet birds through trade across the country. To better understand the modes of transmission of psittacosis to humans, our report highlights the importance of genotyping *C. psittaci* strains in psittacosis patients during diagnosis and the need for future pan-*Chlamydia* detection in pet birds.

## Ethics approval

This study received approval from the Human Research Ethics Committee (EA1912038) and Animal Research Ethics Committee (5264-19) of the University of Hong Kong; as well as the Department of Health [(19-1499) in DH/HT&A/8/2/3 Pt. 3] of the HKSAR Government.

## Consent to participate

Not applicable.

## Consent for publication

Not applicable.

Availability of data and materials

The *OmpA* sequences obtained from this study have been deposited in GenBank with the accession numbers OP594252-OP594256.

## Author Contributions

Methodology, JCKK and ESKP; Investigation, JCKK, ESKP, YWYC; Formal analysis, JCKK and YWYC; Writing –Original Draft, JCKK; Writing – Review & Editing, ESKP and SYYS; Resources, NW and JLLG; Conceptualization, LLMP and SYYS; Supervision, SYYS; All authors read and approved the final manuscript.

## Acknowledgements

We thank the Agricultural, Fisheries, and Conservation Department for communication in facilitating sample collection from pet shops. We are grateful for the help provided by current lab members and alumni for their contribution to this study, including but not limited to Bosco Yuen, Alex Chan, Derek Lam, Pei-Yu Huang, David Chan, Stella Huynh, Yiu Siu, Christy Hung, Verna Shiu, and Joyce Lam. We are also thankful to all participating pet shops and bird owners for their samples and information.

## References

ADM Capital Foundation. 2022. *Wild, threatened, farmed: Hong Kong’s Invisible Pets*. https://www.admcf.org/wp-content/uploads/2022/07/0714_WTF-Report-V03.pdf. Accessed 13 Feb 2024.

Abd El-Ghany, W.A. 2020. Avian Chlamydiosis: A World-wide Emerging and Public Health Threat. Advances in Animal and Veterinary Sciences 8(s2): 82–97. 10.17582/journal.aavs/2020/8.s2.82.97.

Agriculture, Fisheries and Conservation Department. 2024. Permit Terms for Importation/Transhipment of Pet Birds. https://www.afcd.gov.hk/tc_chi/quarantine/qua_ie/qua_ie_ipab/qua_ie_ipab_ibpo/files/permit_terms_for_import_transit_pet_birds_jun11b.pdf. Accessed 13 Feb 2024.

Andersen, A.A. National Animal Disease Center, Respiratory Diseases of Livestock Research Unit), D. Vanrompay. 2000. Avian chlamydiosis [*Chlamydia, Chlamydophila*]. Revue scientifique et technique (International Office of Epizootics) 19: 396–404. 10.20506/rst.19.2.1223.

Anderson, I.E., S.I.F. Baxter, S. Dunbar, A.G. Rae, H.L. Philips, M.J. Clarkson, and A.J. Herring. 1996. Analyses of the Genomes of Chlamydial Isolates from Ruminants and Pigs Support the Adoption of the New Species *Chlamydia pecorum*. International journal of systematic bacteriology 46: 245–251. 10.1099/00207713-46-1-245.

Balsamo, G., A.M. Maxted, J.W. Midla, J.M. Murphy, R. Wohrle, T.M. Edling, P.H. Fish, K. Flammer, D. Hyde, P.K. Kutty, et al. 2017. Compendium of Measures to Control Chlamydia psittaci Infection Among Humans (Psittacosis) and Pet Birds (Avian Chlamydiosis), 2017. *Journal of avian medicine and surgery* 31: 262-282. 10.1647/217-265.

Beeckman, D.S.A., D.C.G. Vanrompay. 2009. Zoonotic *Chlamydophila psittaci* infections from a clinical perspective. Clinical Microbiology and Infection 15: 11–17. 10.1111/j.1469-0691.2008.02669.x.

Billington, S. 2005. Clinical and zoonotic aspects of psittacosis. In practice (London 1979) 27: 256–258. 10.1136/inpract.27.5.256.

Branley, J., N.L. Bachmann, M. Jelocnik, G.S.A. Myers, and A. Polkinghorne. 2016. Australian human and parrot *Chlamydia psittaci* strains cluster within the highly virulent 6BC clade of this important zoonotic pathogen. Scientific Reports 6: 30019. 10.1038/srep30019.

Butaye, P., R. Ducatelle, P. De Backer, H. Vermeersch, J.P. Remon, and F. Haesbrouck. 1997. In vitro activities of doxycycline and enrofloxacin against European *Chlamydia psittaci* strains from turkeys. Antimicrobial Agents and Chemotherapy 41: 2800–2801. 10.1128/AAC.41.12.2800.

Chahota, R., H. Ogawa, Y. Mitsuhashi, K. Ohya, T. Yamaguchi, and H. Fukushi. 2006. Genetic Diversity and Epizootiology of *Chlamydophila psittaci* Prevalent among the Captive and Feral Avian Species Based on VD2 Region of ompA Gene. Microbiology and immunology 50: 663–678. 10.1111/j.1348-0421.2006.tb03839.x.

Chan, D.T.C., E.S.K. Poon, A.T.C. Wong, and S.Y.W. Sin. 2021. Global trade in parrots – Influential factors of trade and implications for conservation. Global ecology and conservation 30: e01784. 10.1016/j.gecco.2021.e01784.

CHP. 2020. Statistics: Number of notifiable infectious diseases by month. https://www.chp.gov.hk/en/static/24012.html. Accessed 13 Feb 2024.

Cong, W., S.Y. Huang, X.X. Zhang, D.H. Zhou, M.J. Xu, Q. Zhao, A.D. Qian, and X.Q. Zhu. 2014. *Chlamydia psittaci* exposure in pet birds. Journal of medical microbiology 63: 578–581. 10.1099/jmm.0.070003-0.

De Freitas Raso, T., G.H.F. Seixas, N.M.R. Guedes, and A.A. Pinto. 2006. *Chlamydophila psittaci* in free-living Blue-fronted Amazon parrots (*Amazona aestiva*) and Hyacinth macaws (*Anodorhynchus hyacinthinus*) in the Pantanal of Mato Grosso do Sul, Brazil. Veterinary microbiology 117: 235–241. 10.1016/j.vetmic.2006.06.025.

Failing, K., P. Theis, and E.F. Kaleta. 2006. Determination of the inhibitory concentration 50% (IC50) of four selected drugs (chlortetracycline, doxycycline, enrofloxacin and difloxacin) that reduce in vitro the multiplication of *Chlamydophila psittaci*. DTW. Deutsche tierarztliche Wochenschrift 113: 412–417.

Feng, Y., Y. Feng, Z. Zhang, S. Wu, D. Zhong, and C. Liu. 2016. Prevalence and genotype of *Chlamydia psittaci* in faecal samples of birds from zoos and pet markets in Kunming, Yunnan, China. Journal of Zhejiang University. B. Science 17: 311–316. 10.1631/jzus.B1500091.

Fudge, A.M. 1997. A Review of Methods to Detect *Chlamydia psittaci* in Avian Patients. Journal of avian medicine and surgery 11: 153–165.

Harkinezhad, T., T. Geens, and D. Vanrompay. 2009. *Chlamydophila psittaci* infections in birds: A review with emphasis on zoonotic consequences. Veterinary microbiology 135: 68–77. 10.1016/j.vetmic.2008.09.046.

Harkinezhad, T., K. Verminnen, C. Van Droogenbroeck, and D. Vanrompay. 2007. *Chlamydophila psittaci* genotype E/B transmission from African grey parrots to humans. Journal of medical microbiology 56: 1097–1100. 10.1099/jmm.0.47157-0.

Hasegawa, M., H. Kishino, and T. Yano. 1985. Dating of the human-ape splitting by a molecular clock of mitochondrial DNA. Journal of molecular evolution 22: 160–174. 10.1007/BF02101694.

Heddema, E.R., E.J. van Hannen, M. Bongaerts, F. Dijkstra, R.J. ten Hove, B. de Wever, and D. Vanrompay. 2015. Typing of *Chlamydia psittaci* to monitor epidemiology of psittacosis and aid disease control in the Netherlands, 2008 to 2013. Euro surveillance : bulletin européen sur les maladies transmissibles 20: 28–35. 10.2807/1560-7917.ES2015.20.5.21026.

Hoang, D.T., O. Chernomor, A. von Haeseler, B.Q. Minh, and L.S. Vinh. 2018. UFBoot2: Improving the Ultrafast Bootstrap Approximation. Molecular Biology and Evolution 35: 518–522. 10.1093/molbev/msx281.

Hogerwerf, L., I. Roof, M.J.K. de Jong, F. Dijkstra, and W. van der Hoek. 2020. Animal sources for zoonotic transmission of psittacosis: a systematic review. BMC Infectious Diseases 20: 192. 10.1186/s12879-020-4918-y.

Inchuai, R., S. Weerakun, H.N. Nguyen, and P. Sukon. 2020. Global Prevalence of Chlamydial Infections in Reptiles: A Systematic Review and Meta-Analysis. *Vector borne and zoonotic diseases (Larchmont*, N.Y.) 21: 32–39. 10.1089/vbz.2020.2654.

Kalyaanamoorthy, S., B.Q. Minh, T.K.F. Wong, A. von Haeseler, and L.S. Jermiin. 2017. ModelFinder: fast model selection for accurate phylogenetic estimates. Nature methods 14: 587–589. 10.1038/nmeth.4285.

Kozuki, E., Y. Arima, T. Matsui, Y. Sanada, S. Ando, T. Sunagawa, and K. Oishi. 2020. Human psittacosis in Japan: notification trends and differences in infection source and age distribution by gender, 2007 to 2016. Annals of epidemiology 44: 60–63. 10.1016/j.annepidem.2020.03.001.

Lang, Z., J. Reiczigel. 2014. Confidence limits for prevalence of disease adjusted for estimated sensitivity and specificity. Preventive veterinary medicine 113: 13–22. 10.1016/j.prevetmed.2013.09.015.

Lee, H., O. Lee, S. Kang, Y. Yeo, J. Jeong, Y. Kwon, and M. Kang. 2023. Prevalence of asymptomatic infections of *Chlamydia psittaci* in psittacine birds in Korea. Zoonoses and public health 70: 451–458. 10.1111/zph.13039.

Lindenstruth, H., J.W. Frost. 1993. Enrofloxacin (Baytril)--an alternative for psittacosis prevention and therapy in imported psittacines. DTW. Deutsche tierarztliche Wochenschrift 100: 364–368.

Liu, C.J., L. Jin and Y. Feng. 2008. Nucleotide accession No. EU856035.1. Chlamydophila psittaci strain SP15 MOMP gene, partial cds. https://www.ncbi.nlm.nih.gov/nuccore/EU856035.1. Accessed 01 May 2023.

Liu, S., K. Li, M. Hsieh, P. Chang, J. Shien, and S. Ou. 2019. Prevalence and Genotyping of *Chlamydia psittaci* from Domestic Waterfowl, Companion Birds, and Wild Birds in Taiwan. *Vector borne and zoonotic diseases (Larchmont*, N.Y.) 19: 666–673. 10.1089/vbz.2018.2403.

Madani, S. A., S.M. Peighambari. 2013. PCR-based diagnosis, molecular characterization and detection of atypical strains of avian *Chlamydia psittaci* in companion and wild birds. Avian pathology 42: 38–44. 10.1080/03079457.2012.757288.

Mina, A., A. Fatemeh, and R. Jamshid. 2019. Detection of *Chlamydia psittaci* Genotypes Among Birds in Northeast Iran. Journal of avian medicine and surgery 33: 22–28. 10.1647/2017-334.

Opota, O., K. Jaton, J. Branley, D. Vanrompay, V. Erard, N. Borel, D. Longbottom, and G. Greub. 2015. Improving the molecular diagnosis of *Chlamydia psittaci* and *Chlamydia abortus* infection with a species-specific duplex real-time PCR. Journal of medical microbiology 64: 1174–1185. 10.1099/jmm.0.000139.

Poon, W.T. 2018. Investigation of prevalence of unregulated trade and the attitude of pet owners in sustainable parrot trade. The University of Hong Kong. 10.5353/th_991044071095403414.

Radomski, N., R. Einenkel, A. Müller, and M.R. Knittler. 2016. *Chlamydia*–host cell interaction not only from a bird’s eye view: some lessons from *Chlamydia psittaci*. FEBS letters 590: 3920–3940. 10.1002/1873-3468.12295.

Ruiz-Laiton, A., N. Molano-Ayala, S. García-Castiblanco, A.M. Puentes-Orozco, A.C. Falla, M. Camargo, L. Roa, A. Rodríguez-López, M.A. Patarroyo, and C. Avendaño. 2022. The prevalence of *Chlamydia psittaci* in confiscated Psittacidae in Colombia. Preventive veterinary medicine 200: 105591. 10.1016/j.prevetmed.2022.105591.

Sachse, K., K. Laroucau, K. Riege, S. Wehner, M. Dilcher, H.H. Creasy, M. Weidmann, G. Myers, F. Vorimore, N. Vicari, et al. 2014. Evidence for the existence of two new members of the family Chlamydiaceae and proposal of *Chlamydia avium* sp. nov. and *Chlamydia gallinacea* sp. nov. Systematic and Applied Microbiology 37: 79–88. 10.1016/j.syapm.2013.12.004.

Shi, Y., J. Chen, X. Shi, J. Hu, H. Li, X. Li, Y. Wang, and B. Wu. 2021. A case of *Chlamydia psittaci* caused severe pneumonia and meningitis diagnosed by metagenome next-generation sequencing and clinical analysis: a case report and literature review. BMC infectious diseases 21: 1–621. 10.1186/s12879-021-06205-5.

Smith, K.A., K.K. Bradley, M.G. Stobierski, and L.A. Tengelsen. 2005. Compendium of measures to control *Chlamydophila psittaci* (formerly *Chlamydia psittaci*) infection among humans (psittacosis) and pet birds, 2005. Journal of the American Veterinary Medical Association 226: 532–539. 10.2460/javma.2005.226.532.

Sukon, P., N.H. Nam, P. Kittipreeya, A. Sara-in, P. Wawilai, R. Inchuai, and S. Weerakhun. 2021. Global prevalence of chlamydial infections in birds: A systematic review and meta-analysis. Preventive veterinary medicine 192: 105370. 10.1016/j.prevetmed.2021.105370.

Tolba, H.M.N., R.M.M. Abou Elez, and I. Elsohaby. 2019. Risk factors associated with *Chlamydia psittaci* infections in psittacine birds and bird handlers. Journal of applied microbiology 126: 402–410. 10.1111/jam.14136.

Trifinopoulos, J., L. Nguyen, A. von Haeseler, and B.Q. Minh. 2016. W-IQ-TREE: a fast online phylogenetic tool for maximum likelihood analysis. Nucleic Acids Research 44: W232–W235. 10.1093/nar/gkw256.

Vanrompay, D. 2019. Avian chlamydiosis. In Diseases of Poultry. Edited by David E. Swayne, Martine Boulianne, Catherine M. Logue, et al. New Jersey. John Wiley & Sons, Inc. Chapter 24.

Vilela, D.A.R., S.Y. Marin, M. Resende, H.L.G. Coelho, J.S. Resende, F.C. Ferreira-Junior, M.C. Ortiz, A.V. Araujo, T.F. Raso, and N.R.S. Martins. 2020. Phylogenetic analyses of *Chlamydia psittaci* OmpA gene sequences from captive blue-fronted Amazon parrots (*Amazona aestiva*) with hepatic disease in Brazil. Revue scientifique et technique (International Office of Epizootics) 38: 711–719. 10.20506/rst.38.3.3020.

Yin L., I.D. Kalmar, J. Boden, D. Vanrompay. Chlamydia psittaci infections in Chinese poultry: a literature review. 2015, 71(3):473–482. 10.1017/S0043933915002226.

Zhang, N.Z., X.X. Zhang, D.H. Zhou, S.Y. Huang, W.P. Tian, Y.C. Yang, Q. Zhao, and X.Q. Zhu. 2015. Seroprevalence and genotype of *Chlamydia* in pet parrots in China. Epidemiology and infection 143: 55–61. 10.1017/S0950268814000363.

Zhang, Z., H. Zhou, H. Cao, J. Ji, R. Zhang, W. Li, H. Guo, L. Chen, C. Ma, M. Cui, et al. 2022. Human-to-human transmission of *Chlamydia psittaci* in China, 2020: an epidemiological and aetiological investigation. The Lancet. Microbe 3: e512–e520. 10.1016/S2666-5247(22)00064-7.

